# Clockwise and counterclockwise hysteresis characterize state changes in the same aquatic ecosystem

**DOI:** 10.1101/2020.05.01.073239

**Authors:** Amanda C. Northrop, Vanessa Avalone, Aaron M. Ellison, Bryan A. Ballif, Nicholas J. Gotelli

## Abstract

Incremental increases in a driver variable, such as nutrients or detritus, can trigger abrupt shifts in aquatic ecosystems. Once these ecosystems change state, a simple reduction in the driver variable may not return them to their original state. Because of the long time scales involved, we still have a poor understanding of the dynamics of ecosystem recovery after a state change. A model system for understanding ecosystem recovery is the aquatic microecosystem that inhabits the cup-shaped leaves of the pitcher plant *Sarracenia purpurea*. With enrichment of organic matter, this system flips within 1 to 3 days from an oxygen-rich state to an oxygen-poor (hypoxic) state. In a replicated greenhouse experiment, we enriched pitcher plant leaves at different rates with bovine serum albumin (BSA), a molecular substitute for detritus. Changes in dissolved oxygen ([O_2_]) and undigested BSA concentration were monitored during enrichment and recovery phases. At low enrichment rates, ecosystems showed a substantial lag in the recovery of [O_2_] (clockwise hysteresis). At intermediate enrichment rates, [O_2_] tracked the levels of undigested BSA with the same profile during the enrichment and recovery phases (no hysteresis). At high enrichment rates, we observed a novel response: changes in [O_2_] were proportionally larger during the recovery phase than during the enrichment phase (counter-clockwise hysteresis). These experiments demonstrate that detrital enrichment rate can modulate a diversity of hysteretic responses in a single aquatic ecosystem. With counter-clockwise hysteresis, rapid reduction of a driver variable following high enrichment rates may be a viable restoration strategy.

---

Anthropogenically enriched ecosystems often exhibit complex dynamics and regime shifts^1^. Such shifts occur when incremental changes in a driver variable suddenly tip these systems from one basin of attraction to another^1–5^. Early studies emphasized the importance of forecasting impending regime shifts by signature statistical changes in the autocorrelation^6^ or the variance^7^ of a response variable. However, the lead times^8^ and sampling frequencies^9^ necessary to detect early warning signals are usually too long to implement for practical management strategies. Equally important for management is how a system recovers after a collapse. Aquatic ecosystems that collapse rapidly often recover slowly^10–12^, and may remain in an altered state long after enrichment has ceased. A general mechanism that might cause such a lag is hysteresis – a phenomenon in which the relationship between a response variable and driver variable depends on the state of the system^13^. In hysteretic systems, changes in the response variable lag behind those in the driver variable due to feedback loops between the response variable and other components of the system.

In spite of their potential importance to management and restoration ecology^14^, hysteretic responses of recovering enriched ecosystems have rarely been fully characterized. The limitations are the long time frame over which recovery occurs^15^ relative to the generation times of the component species^16^, and the challenge of replicating whole-ecosystem measurements in both experimental and correlative studies^13, 17^. Here we conducted a greenhouse experiment using a model aquatic ecosystem that can collapse within 1 to 3 days of detrital enrichment and recover in 10 to 35 days. This process occurs over many generations of the bacteria driving ecosystem collapse and recovery. We characterized the recovery trajectory in replicated ecosystems receiving low, intermediate, and high levels of organic matter enrichment, and discovered a range of recovery dynamics, including a novel counter-clockwise hysteretic loop in response to high levels of enrichment.

The pitchers of *S. purpurea* are cup-shaped leaves that fill with rainwater and capture arthropod prey^18^, which forms the base of a “brown” detrital food web^19^. The drowned prey initially are shredded by aquatic larvae of sarcophagid flies and midges, but the complete breakdown and mineralization of the prey is predominantly the result of microbial activity^19^. The mineralized nutrients are quickly assimilated and translocated to plant tissues^19^. In the field, less than 1% of prey encounters result in successful capture^20, 21^. With low detrital inputs and active photosynthesis by the plant, this aquatic microecosystem is normally in an oligotrophic state, with low prey abundance and [O_2_] close to 20%. But with excessive loading of prey or detritus, [O_2_] collapses within 8 hours to less than 5%^22^. With no additional detrital enrichment, the system persists in this anoxic eutrophic state for many days or weeks until the excess prey is transformed and [O_2_] slowly recovers^22^. Except for the compressed time scale, this trajectory of enrichment, sudden collapse of [O_2_], and slow recovery characterizes both photosynthetic “green” food webs that experience direct nutrient enrichment and tightly-linked “brown” food webs enriched with organic matter from the decomposition of primary-producer biomass^23^.

In this study, we used bovine serum albumin (BSA) as a molecular substitute for arthropod prey. We created a BSA cocktail that has the same nutrient content and stoichiometric ratios as natural insect prey^24^. Because BSA is water-soluble, we could easily monitor the concentration of unprocessed BSA with a non-destructive Bradford assay^25^. Pilot experiments confirmed that collapse in [O_2_] after BSA addition mimics collapse after arthropod prey addition (Supplementary Data Fig. 1) and that BSA is stable in an aqueous solution and does not break down in the absence of microbial activity (Supplementary Data Fig. 2).

**Figure 1.**
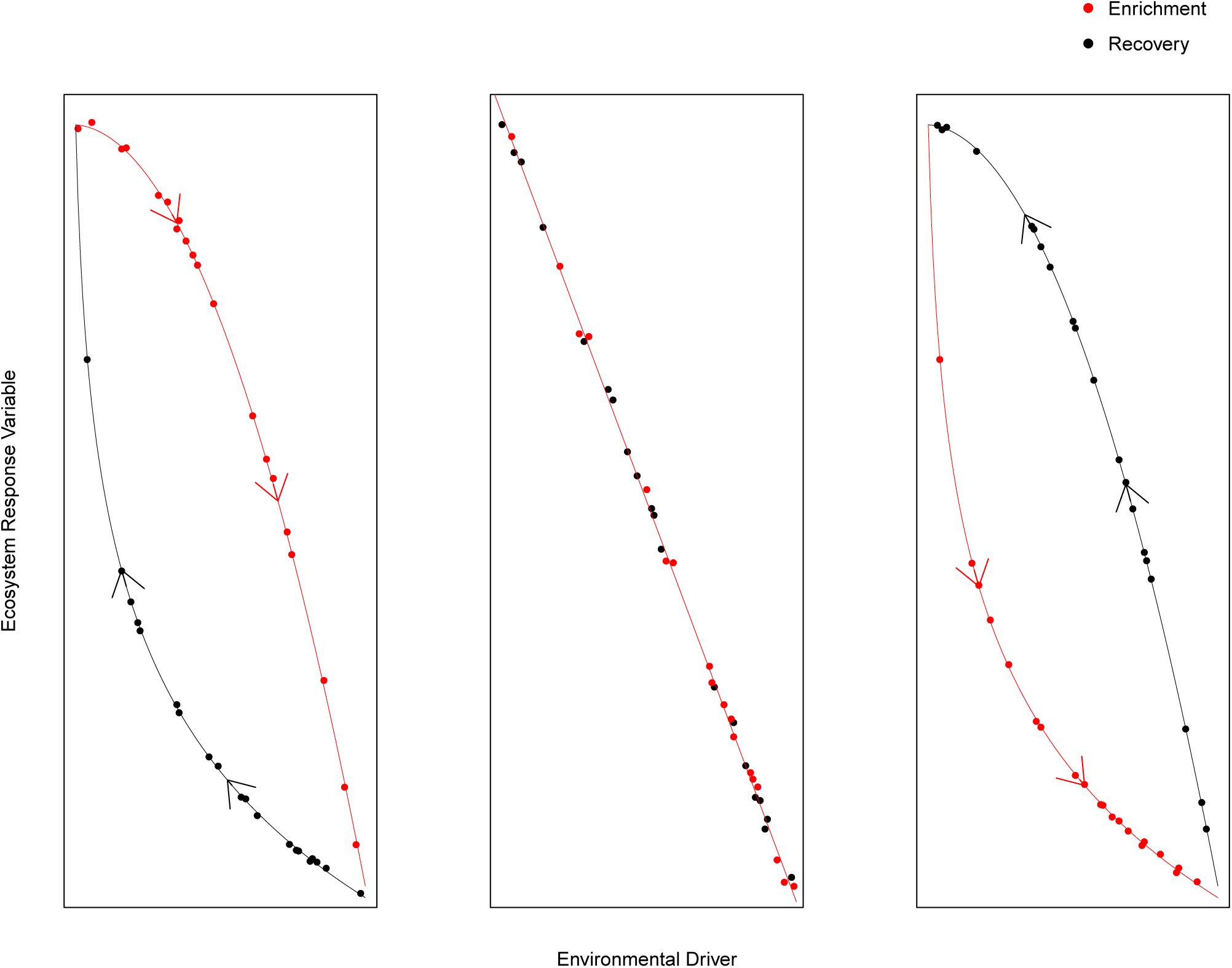
Possible hysteresis loops in driver-response relationships. Clockwise hysteresis (left panel) (*HI*_MEAN_ *>* 0), counterclockwise hysteresis (right panel) (*HI*_MEAN_ < 0), and environmental tracking (center panel) (*HI*_MEAN_ = 0).

**Figure 2.**
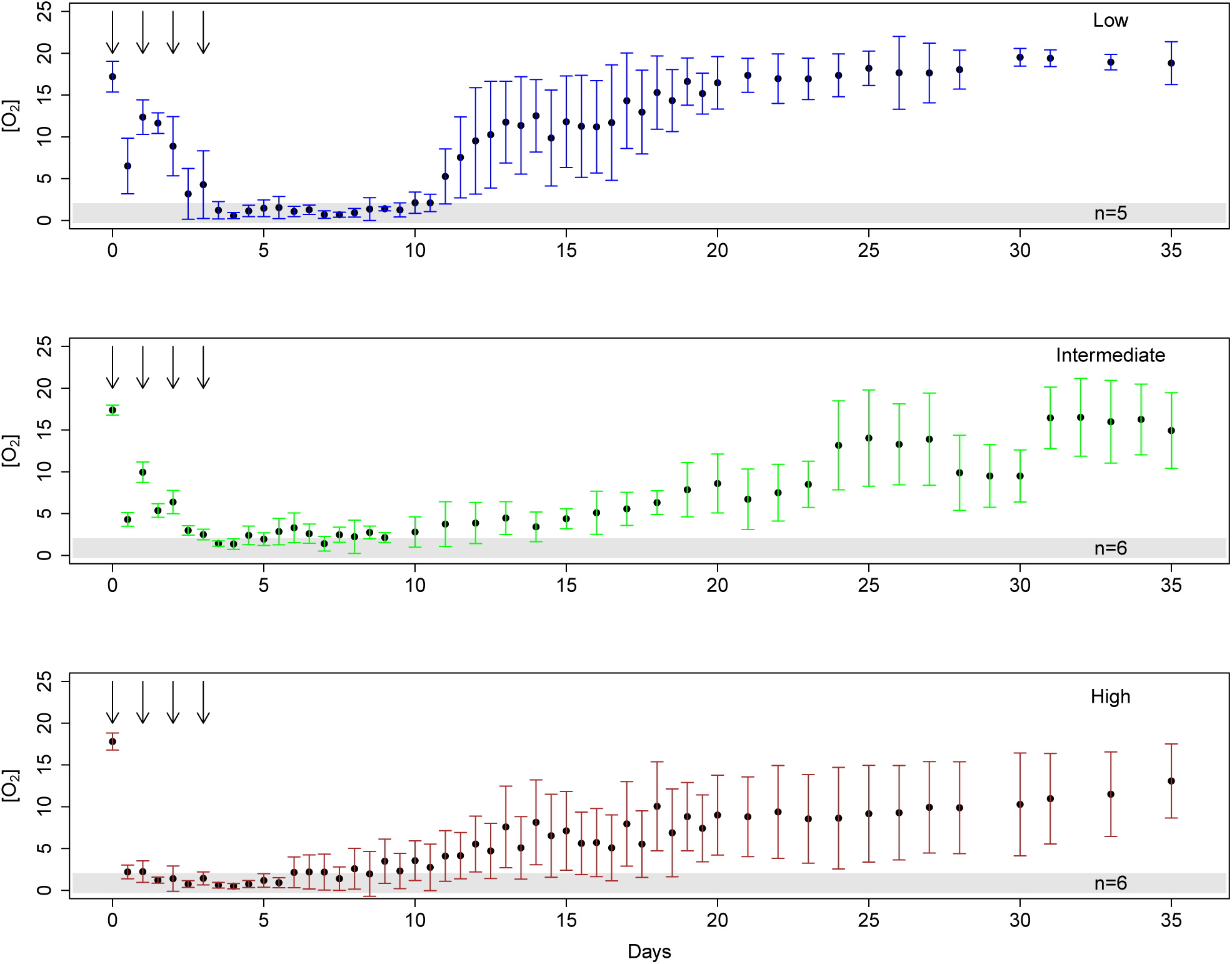
The addition of bovine serum alubmin (BSA) to replicate pitchers causes hypoxia at all levels of enrichment greater than or equal to 0.05 mg BSA/ml pitcher fluid. Time series of average dissolved oxygen concentration (expressed as a percent) from all replicate pitchers in low, intermediate and high BSA addition treatments. Error bars represent one standard deviation of the mean. Vertical arrows represent the timing of BSA additions. Gray bands denote the range of [O_2_] considered hypoxic (<2%).

We ran a greenhouse experiment in July and August in 2015 and 2016 at the University of Vermont’s Biological Research Complex. Six replicate pitchers in each treatment were enriched in a press experiment with low (0.5 mg/mL), intermediate (2.0 mg/mL), or high (5.0 mg/mL) concentrations of BSA cocktail. [O_2_] was monitored and pitcher fluid sampled twice daily during enrichment. Once mean [O_2_] was less than 2.0%, BSA enrichment was halted. Pitcher fluid collection and [O_2_] monitoring continued until [O_2_] in all replicate pitchers was within 2% of [O_2_] in matched control pitchers of similar volume. [O_2_] in all replicate pitchers measured between 15.6% and 20.1% (mean = 17.2%) on Day 0. The amount of unprocessed BSA in all pitcher fluid samples was measured using a Bradford Assay^25^.

We first plotted [O_2_] against BSA concentration for each of the enrichment and recovery trajectories in all replicates and for replicates pooled within treatments. We then calculated mean hysteresis indices for each treatment, using a common metric for river-discharge hysteresis (HI_MEAN_)^26^, to determine the direction and strength of hysteresis at low, intermediate, and high levels of BSA enrichment. Positive values of (HI_MEAN_) denote a clockwise hysteresis loop and negative values indicate a counterclockwise loop (Fig 1). The magnitude of the index (from −1 to +1) indicates the strength of hysteresis, with zero indicating no hysteresis^26^.

## Results

Within 5 d after the start of enrichment, [O_2_] in all but one enriched pitcher was ≤ 2% (Fig.2). The average time to collapse differed significantly among the three enrichment treatments (*F*_2,14_ = 14.38, *p* = 0.0004) for low 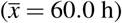, intermediate 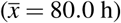, and high 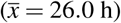 enrichment treatments. After enrichment was halted, [O_2_] recovered gradually for another 30 d (Fig.2). Variation in mean [O_2_] near the end of the experiment was due to the failure of pitchers in the intermediate (4 pitchers) and high (5 pitchers) BSA treatments to reach control [O_2_] levels by Day 35 (*χ*^2^ = 8.30, df=2, p=0.016; Fig.2, Supplementary Data Fig. 3). Using the terminal date of the experiment for pitchers that did not fully return to control [O_2_] levels (a conservative estimate), there was a significant difference (*F*_2,14_ = 18.67, *p* = 0.0001) among the treatments in recovery time 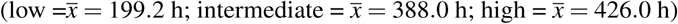. In all treatments, there was a substantial reservoir of undigested BSA remaining in pitchers after the cessation of enrichment that persisted for weeks (Supplementary Data Fig. 4).

**Figure 3.**
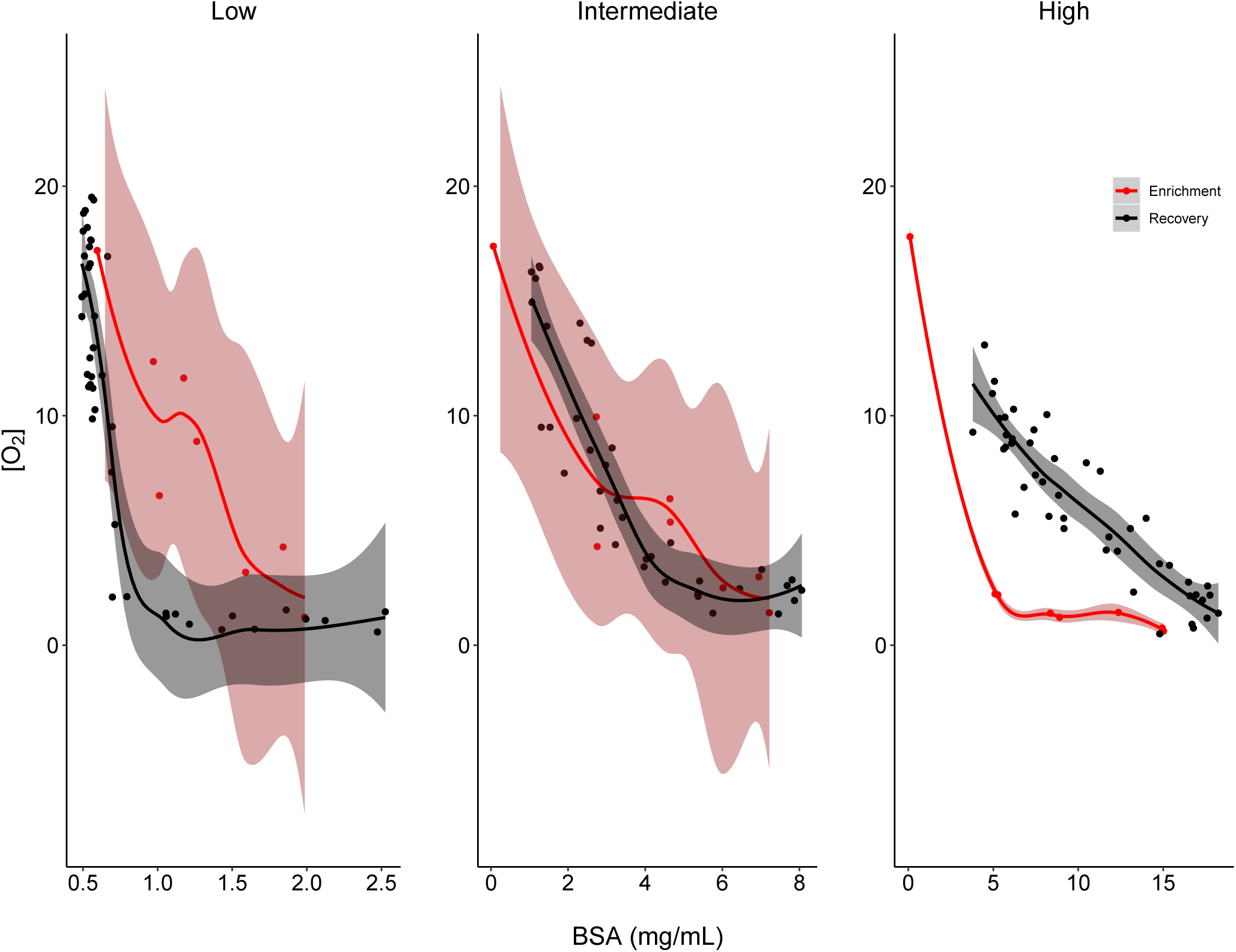
Altering a single driver variable, bovine serum albumin (BSA), elicits counterclockwise and clockwise hysteresis in an aquatic ecosystem. Local regression (loess) curves (span = 0.5) fitted to state-space plots for all replicates in low, intermediate, and high BSA addition treatments with 95% confidence intervals. Red and black points, lines, and shading denote data from the enrichment and recovery phases, respectively.

**Figure 4.**
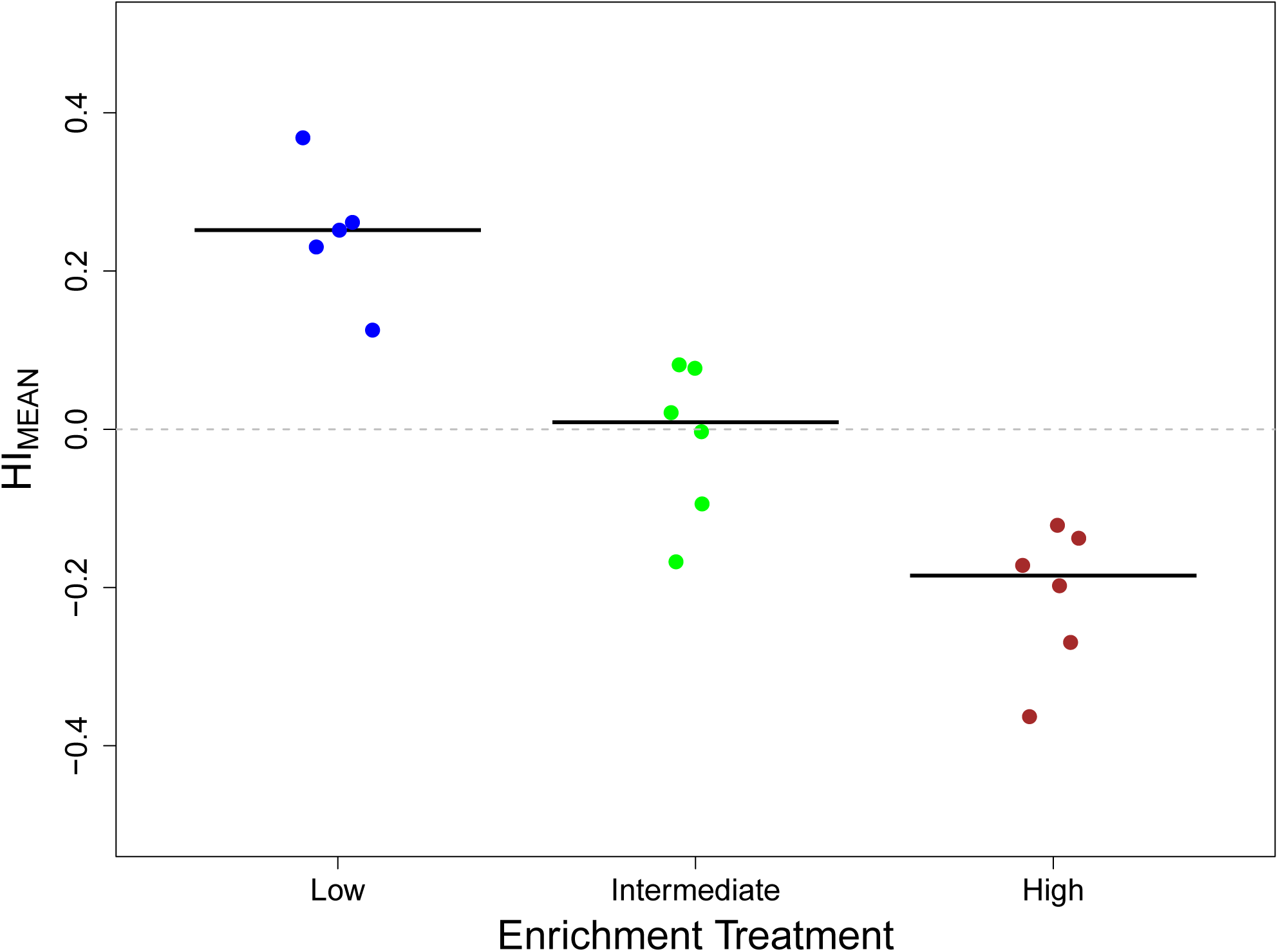
The strength and direction of hysteresis, indicated by HI_MEAN_, was significantly different between all enrichment treatments (ANOVA, F_2,16_ = 35.26, p < 0.001; Tukey HSD, p < 0.001). HI_MEAN_ describes the magnitude of hysteresis (from 0 to 1, where 0 = no hysteresis) and direction (negative = counterlockwise, positive = clockwise). Blue, green, and red dots represent HI_MEAN_ of replicates experiencing low (*n*=5), intermediate(*n*=8), and high (*n*=6) levels of enrichment, respectively. Solid horizontal lines represent mean HI_MEAN_ values in each treatment and the dotted horizontal line denotes where HI_MEAN_ = 0.

Pitchers receiving a low rate of BSA enrichment displayed a classic clockwise hysteresis loop in the relationship between BSA concentration and [O_2_], with a lag in [O_2_] recovery relative to decreasing BSA concentration during the recovery phase (HI_MEAN_ = 0.247; Fig. 3, Fig. 4, Supplementary Data Fig. 5). Hysteresis was absent at intermediate levels of enrichment: enrichment and recovery curves overlapped, suggesting a responsive tracking of BSA concentration by [O_2_] (HI_MEAN_ = − 0.015; Fig. 3, Supplementary Data Fig. 5). High levels of BSA enrichment yielded a counterclockwise hysteresis loop: [O_2_] changed faster per unit change in BSA concentration during recovery than enrichment (HI_MEAN_ = −0.210; Fig. 3, Supplementary Data Fig. 5).

## Discussion

Here we have shown that an enriched aquatic ecosystem can display a diverse set of hysteretic responses modulated by changes in a single driver variable. We discovered an unexpected counterclockwise hysteresis loop resulting from high levels of enrichment. Counterclockwise hysteresis has been detected in some physical and biological^27–29^ systems, but has rarely been documented in ecological studies. These few ecological studies with counterclockwise hysteresis loops quantify static patterns of hydrological relationships along a spatial gradient, rather than measuring ecosystem dynamics and changes through time^30, 31^.

An analytical systems model of the *S. purpurea* ecosystem predicts clockwise hysteresis as a result of smooth changes in photosynthesis coupled with an abrupt increase in biological oxygen demand (BOD)^32^. Indeed, the addition of organic matter to pitchers causes an abrupt increase in BOD resulting from decomposition^22, 32^ by carbon-limited^33^ bacteria. Pitchers in all enrichment treatments saw an rapid decline in [O_2_] following the initial enrichment phase, suggesting that BOD increased rapidly (Fig. 2). This rapid change in BOD may have contributed to clockwise hysteresis at low levels of enrichment, but doesn’t account for the counterclockwise hysteresis at high enrichment levels. In other systems, clockwise hysteresis is generally the result of positive feedback loops^34^.

Hysteretic ecosystems may require larger restoration efforts to recover from a regime shift^35^. For example, in eutrophic shallow lakes, a simple reduction in phosphorus input does not lead to a proportional recovery in macrophyte cover^10^ or community structure^12^. Further, these communities do not fully recover in the time frames in which they are studied^12^ and may effectively remain permanently degraded. These examples and our work highlight the importance of applying a dynamic regime concept^35^ to ecosystem management and restoration. Such an approach would include testing for hysteresis, characterizing feedbacks that maintain undesirable regimes, and identifying if and how system variables change as a result of a regime shift^36^.

In ecosystems where hysteresis is counter-clockwise, rapid reduction in a driver variable from high to low levels may be a successful restoration strategy. In contrast, systems that have experienced chronic low-levels of enrichment may exhibit clockwise hysteresis that requires more extreme reductions of the driver variable, or alternative restoration strategies^36^, to restore. Although there is still an important need for early warning signals, past histories of high versus low enrichment may dictate different restoration strategies for collapsed ecosystems.

## Methods

### Experimental Setup

Pitcher plants were grown in the University of Vermont’s Biological Research Complex in a temperature-controlled greenhouse. Recently fully-formed pitchers greater than 8 ml in volume were randomly chosen and assigned to control and enrichment treatments. Pitchers were randomly placed on a greenhouse bench to account for spatial variation in greenhouse conditions. All pitchers were rinsed twice with reverse-osmosis water and allowed to dry overnight prior to the start of the experiment. Pitcher fluid was collected from Molly Bog (44.50 °N 72.64 °W), an ombrotrophic bog in Morristown, VT, USA. Fluid was collected randomly from multiple pitchers in the field on July 6, 2015 using a sterile pipette and transported immediately to the greenhouse where it was passed through the 30 micrometer frit bed of a chromatography column (Bio-Rad, Hercules, CA) to remove macrobes. The filtered fluid was mixed evenly and added to experimental pitchers.

### Organic Matter Loading

Pitchers were enriched with either 5.0 (high concentration) or 0.5 (low concentration) mg of organic matter per mL of pitcher fluid. A set of control pitchers received no organic matter. For the first four days of the experiment, experimental pitchers were loaded with organic matter (between 9:00 am and 9:45 am) following pitcher-fluid sampling and [O_2_] measurement. Previous experiments used dried and ground bald-faced hornets (*Dolichovespula maculata*) as organic matter, which has similar elemental composition to the most common prey item of pitchers plants^22, 37^. For this experiment, we created a “BSA cocktail” using cOmniPur bovine serum albumin (BSA) fraction V (VWR) plus trace elements and salts. BSA is a soluble protein that is well-studied, relatively inexpensive, and has been used previously in experiments as a carbon and energy source for microbial communities^32^. In a pilot experiment, enrichment with the BSA cocktail yielded similar [O_2_] profiles to those seen during wasp enrichment.

The BSA cocktail included the trace elements potassium (KCl), calcium (CaCl_2_), sodium (NaCl), magnesium (MgSO_4_), and manganese (MnCl_2_) in the following ratio: 1: 0.115: 0.044: 0.026: 0.0001. This ratio is similar to the elemental composition of royal jelly and adult honeybees^33,34^ and most closely represents trace-element ratios in bald-faced hornets. We also added 1.5 *µ*g of DNA sodium salt from salmon testes (Sigma) per mg of BSA based on DNA yields between 1.4 and 1.5 *µ*g/mg from *Apis melifera*^35^. The BSA cocktail was pre-made for each pitcher, filtered through a sterile 0.2 *µ*m filter, stored in sterile 1-ml microtubes, and frozen at −20 °C until loaded into pitchers. The BSA cocktail was introduced to pitchers using sterile 8-mL transfer pipettes and mixed into pitchers by drawing pitcher fluid in and out of the pipette three times. We were careful to limit the amount of air deposited by and introduced into the transfer pipettes during loadings so that [O_2_] measurements would be minimally affected. Control pitchers received sham loadings in which pitcher fluid was drawn into and deposited from a transfer pipette to account for the effect of mixing. All pitchers were topped off with reverse osmosis water after enrichment/sham loadings.

### Data and Sample Collection

We measured [O_2_] twice a day each day from day 0 to day 20 at 8:30 am and 5:00 pm (±2 hrs), once per day from day 20 to day 28 at 8:30 am (±2 hrs), and once on Days 30, 31, 33, and 35 (8:30 am ±2 hrs). [O_2_] was measured using a D-166MT-1S microelectrode (Lazar Research Laboratories). The microelectrode was calibrated prior to each sampling event according to the manufacturer’s instructions. A single microelectrode was used to take readings from each pitcher and was rinsed twice with reverse osmosis water between readings. The order of readings from different replicates was randomized so that changes in temperature and sunlight over the sampling period were not confounding factors. During readings, the microelectrode was placed 2.5 cm below the surface of the pitcher fluid and swirled so that the more oxygen-rich pitcher fluid at the top of the pitcher was mixed and readings reflected average [O_2_]. Due to the sensitivity of the microelectrode, readings were taken as soon as the reader settled on a value for more than 10 s.

Pitcher fluid was sampled following each DO measurement for spectrophotometric analysis using a Bradford assay^25^. Using sterile pipette tips, a 300 *µ*L aliquot was taken from 2.5 cm below the surface of each pitcher and placed in a sterile 1 mL microfuge tube. Sample tubes were immediately transported to the lab where they were centrifuged at 13,000 × *g* for two minutes. The supernatant containing soluble BSA was removed, placed in a sterile 1 mL microfuge tube, and stored at *−*80 °C until analyzed. [O_2_] was measured and pitcher fluid samples were taken once or twice a day for a total of 36 days.

### BSA Loading Validation via SDS-PAGE

To validate that the majority of protein in extracted pitcher fluid was BSA, we ran a time series of pitcher fluid from two replicates each of the high- and low-concentration loading treatments on gels using SDS-PAGE next to known concentrations of BSA (0.1, 0.5, and 5.0 mg/ml) and a BenchMark Pre-stained Protein Ladder (Invitrogen). Ten *µ*L aliquots of pitcher fluid from days 0, 2, 4, 6, 9, 12, 15, 18, 21, 24, 27, and 30 were added to 90 *µ*L of bromophenol blue sample buffer (150 mM Tris pH 6.8, 2% SDS, 5% beta-mercaptoethanol, 7.8% glycerol) and boiled at 95 °C for five minutes. After centrifugation at 13,000 × *g* for 30 seconds, 10 microliters of each sample was mixed with 10 *µ*L of sample buffer and loaded into separate lanes of a 10% polyacrylamide (37.5:1 acrylamide:bis-acrylamide). Gels were subjected to SDS-PAGE and stained with Coomassie.

### Bradford Assay

We used a Bradford assay^25^ to determine the concentration of BSA. Bradford assays were done using diluted samples after generating a standard curve with known amounts of BSA. Absorbance was measured using a Biophotometer Plus (Eppendorf) at an optical density of 600 nm. To streamline 2016 BSA concentration data collection, assays were conducted on a 96-well plate. Absorbance was measured at 545 nm with a Synergy HTX (Biotek).

### 2016 Experiment

A similar experiment was done in the summer of 2016 using identical field and greenhouse methods. This experiment was started on July 3rd, 2016 in an attempt to control for seasonal variation in the initial microbial community composition and prey capture of field-collected pitcher fluid. The only difference in methods from 2015 to 2016 is that pitchers were enriched with 2 mg/mL BSA/day and pitchers were not removed from the experiment until they had reached control conditions.

### Data Analysis

We used R software (version 3.4.3) to plot the hysteresis loops using the loess function to fit curves (span = 0.5) and 95% confidence intervals to each of the enrichment and recovery trajectories. Hysteresis indices (HI_MEAN_) were calculated for each replicate pitcher to determine the direction and magnitude of hysteresis in the relationship between BSA concentration and [O_2_] using the improved method proposed by Lloyd et al.^26^. Mean HI_MEAN_s were compared among treatments using a one-way analysis of variance (ANOVA) followed by a Tukey’s *post-hoc* test. All statistical analyses were performed in R Studio (v1.1.442, RStudio, Boston, Massachusetts, USA).

### Data availability

All data generated during or analysed during the current study are available from the Harvard Forest Data Archive under ID Number HF334 (doi:<u>10.6073/pasta/ebf6b175a6f6e44d3e9747c13f0d376c</u>).

## Acknowledgements

This work was funded by the National Science Foundation (Grant Numbers 1144055 and 1144056). Research reported in this publication was supported by an Institutional Development Award (IDeA) from the National Institute of General Medical Sciences of the National Institutes of Health under grant number P20GM103449. Its contents are solely the responsibility of the authors and do not necessarily represent the official views of NIGMS or NIH. The authors thank Thomm Buttolph for assistance with the microplate Bradford Assays and Dr. Donna Rizzo for her expertise.

## Author Contributions

A.C.N., N.J.G, A.M.E., and B.A.B conceived the experiment, A.C.N. and V.A. conducted the experiments, A.C.N. analyzed the results. A.C.N. and N.J.E. authored the first draft and all authors contributed to revisions.

## Additional Information

**Supplementary Information** is available for this paper. The authors declare no competing financial interests.

## Supplementary Data Figure Legends

**Supplementary Data Figure 1.**
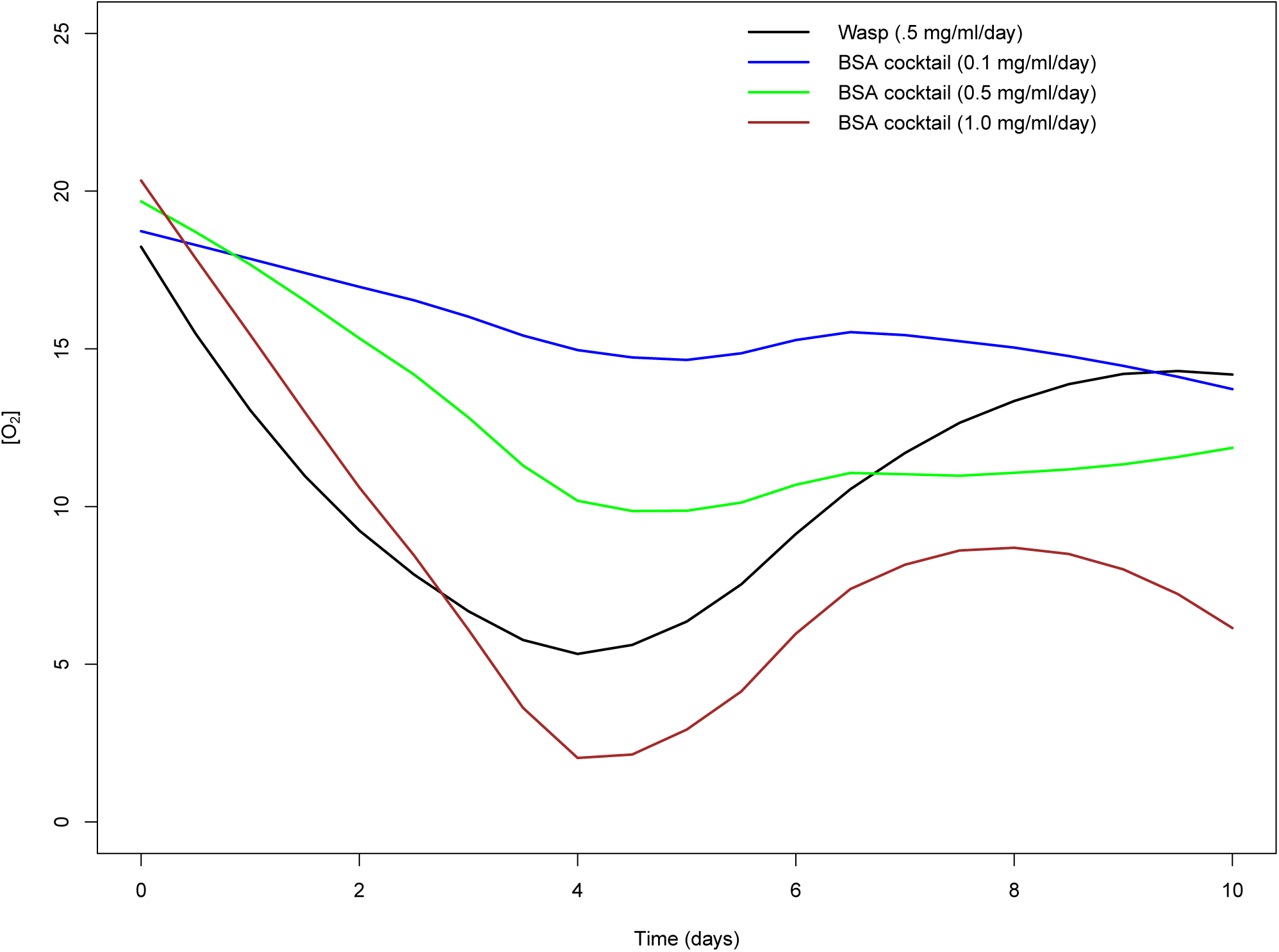
Pitchers enriched with a BSA cocktail experienced decline and recovery of [O_2_] similar to pitchers enriched with ground and autoclaved wasps used in previous experiments. Lines represent a loess fit (span = 0.5) to mean [O_2_] time series for pitchers enriched with 0.5 mg/ml/day ground wasp (black), or 0.1 (blue), 0.5 (green), or 1.0 (brown) mg/ml/day of BSA cocktail over a 10 days during a greenhouse enrichment pilot experiment.

**Supplementary Data Figure 2.**
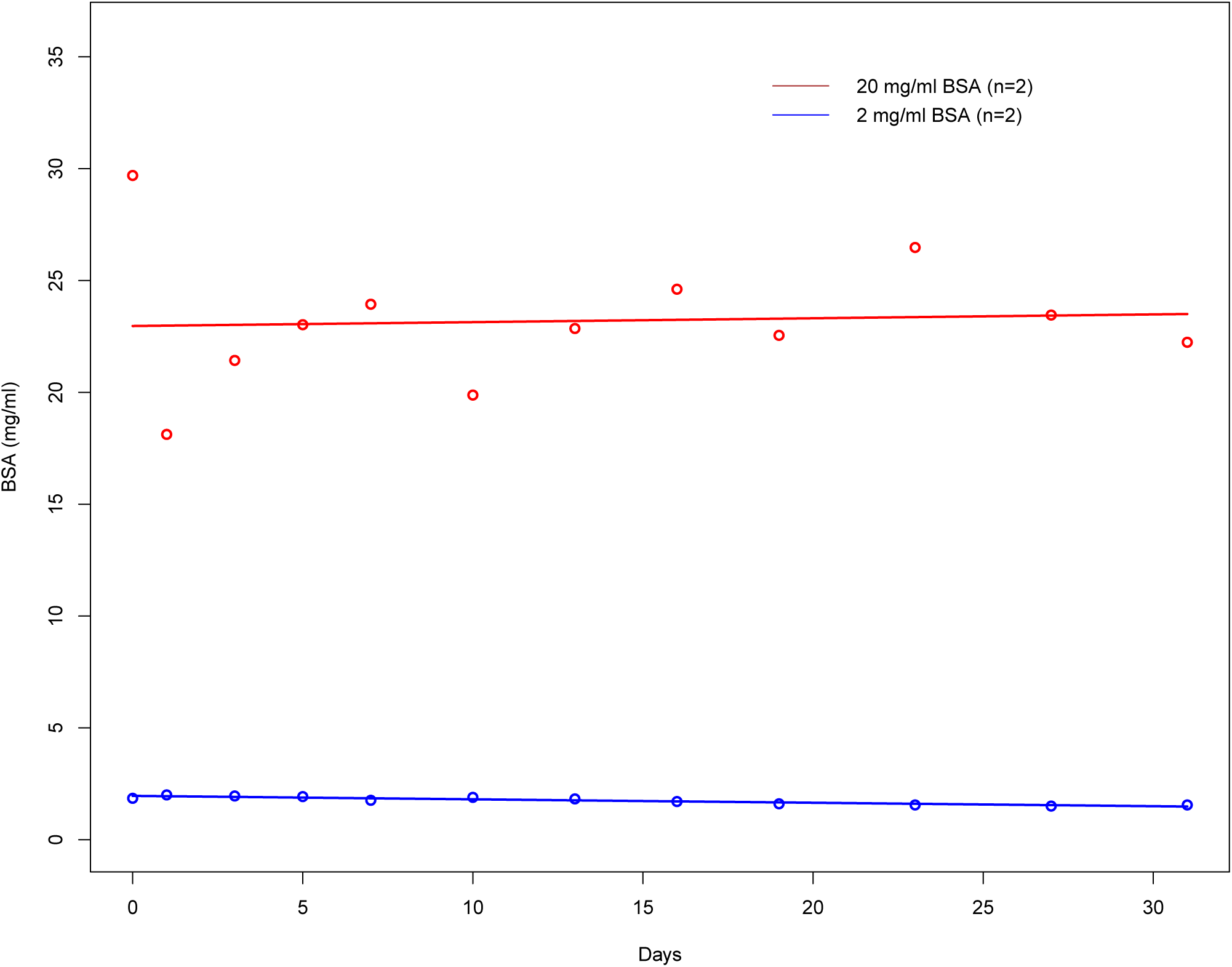
Bovine serum albumin (BSA) does not degrade significantly over the course of 30 days in the absence of microbes. Brown and blue lines represent time series of average BSA concentration for replicates with initial BSA concentrations of 20 mg/mL BSA or 2 mg/mL BSA, respectively.

**Supplementary Data Figure 3.**
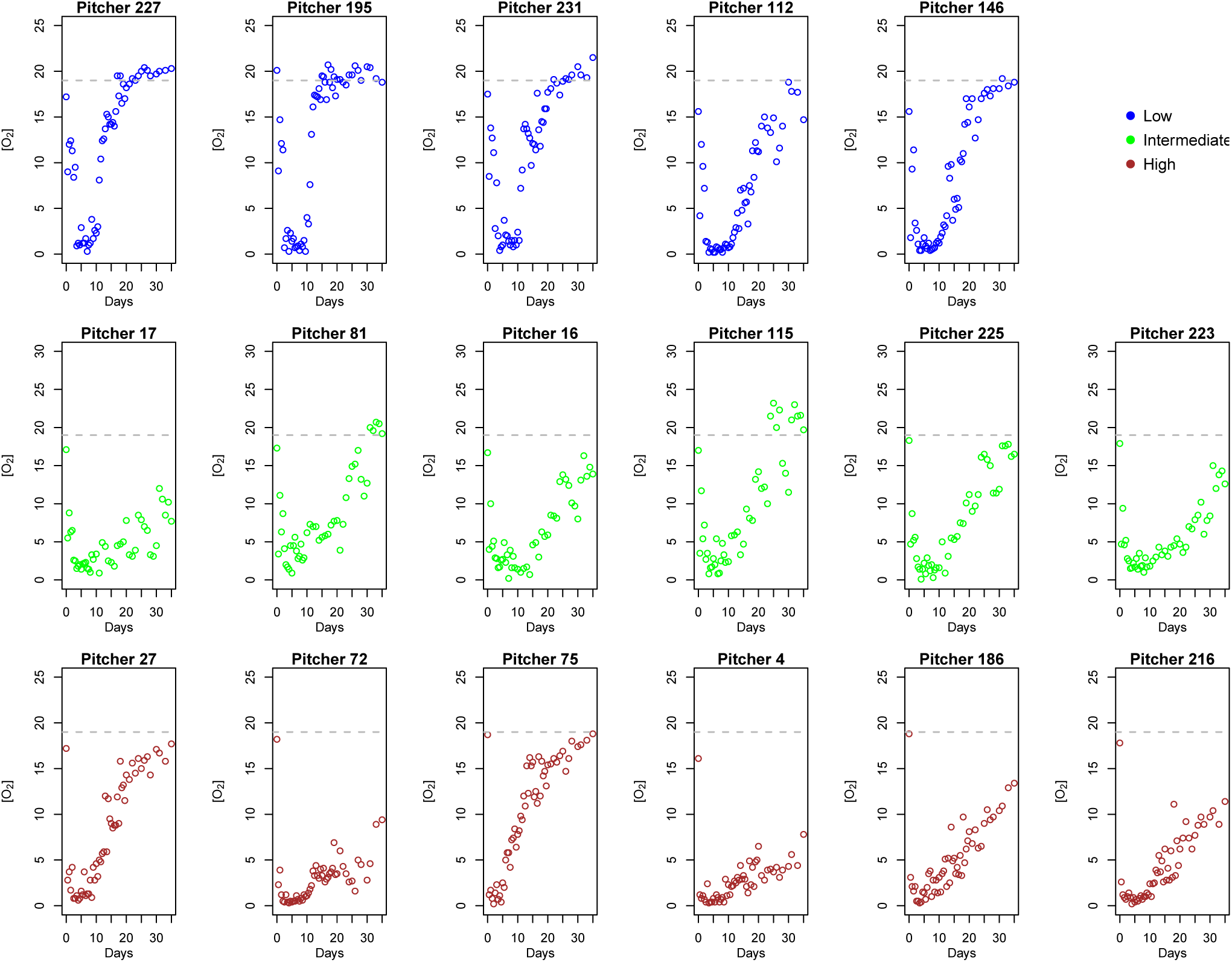
Time series of [O_2_] in replicate pitchers reveals that [O_2_] returned to initial concentrations at the end of 35 days in all low enrichment rate replicates, most intermediate enrichment rate replicates, and half of the high enrichment rate replicates. The gray horizontal dashed lines represent the average initial oxygen concentration on day 0. Blue, green, and brown points represent [O_2_] time series for low, medium, and high enrichment rate replicates, respectively.

**Supplementary Data Figure 4.**
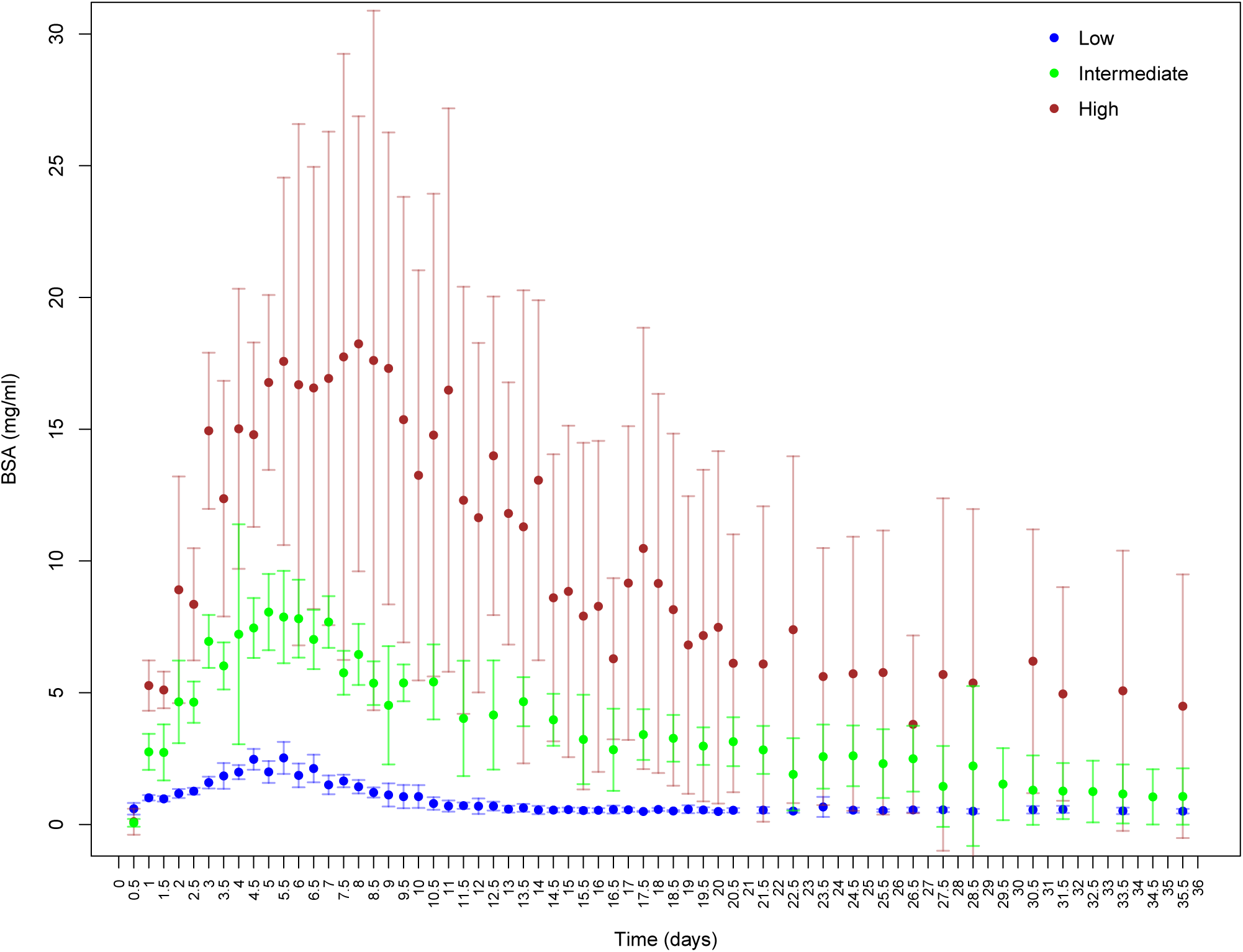
A portion of bovine serum albumin (BSA) remained unprocessed for days to weeks after cessation of enrichment. Time series of average BSA concentration (mg/mL) in low (blue), intermediate (green), and high (brown) enrichment treatments. The vertical dashed line represents the cessation of enrichment on day 4.

**Supplementary Data Figure 5.**
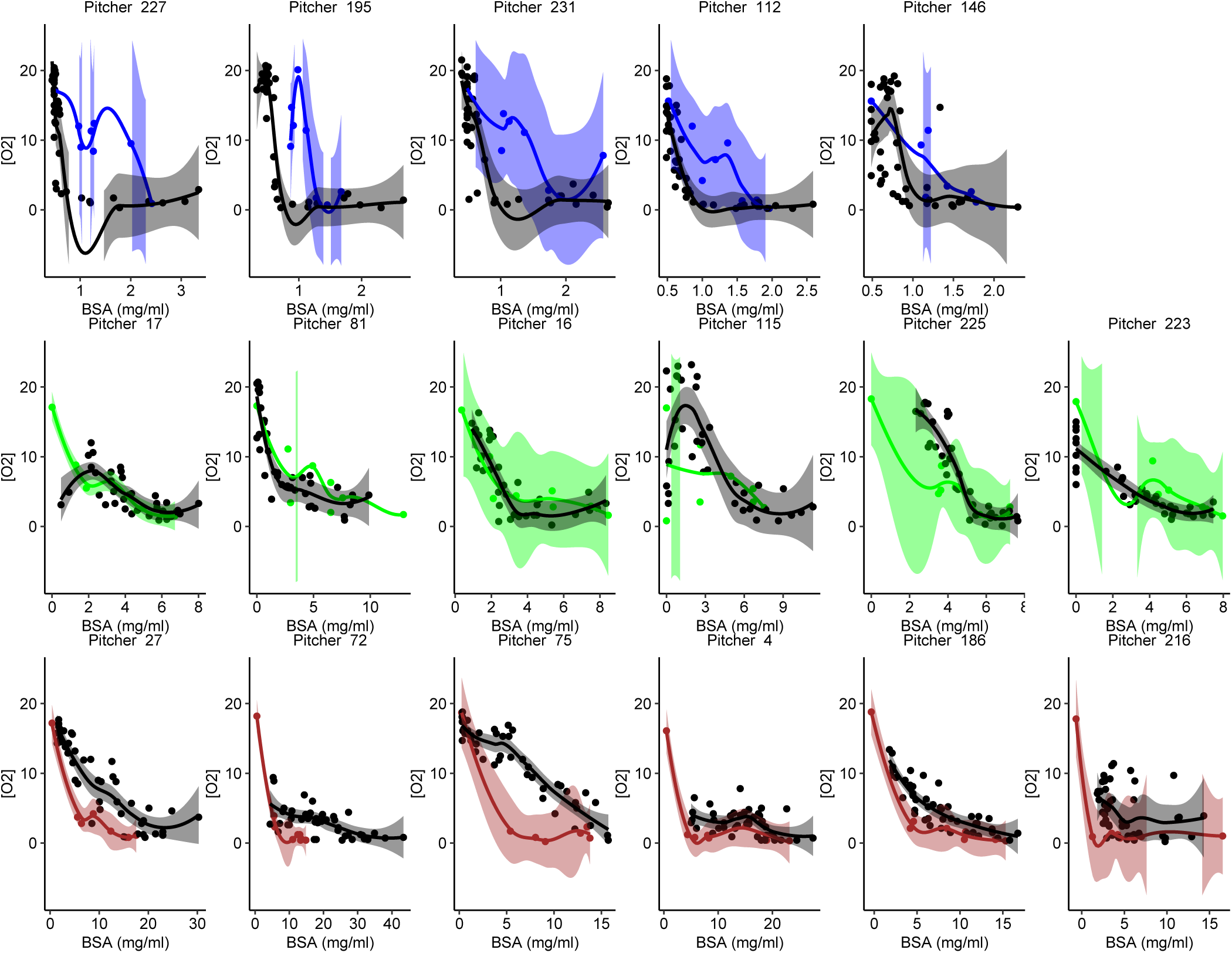
Local regression (loess) curves (span = 0.5) fitted to state-space plots for all replicates in low, intermediate and, high bovine serum albumin (BSA) enrichment treatments with 95% confidence intervals. Blue, green, and brown points, lines, and shading denote data from low, intermediate, and high enrichment phases, respectively. Black points, lines, and shading denote data from the recovery phase in each replicate.

